# Antagonists of the stress and opioid systems restore the functional connectivity of the prefrontal cortex during alcohol withdrawal through divergent mechanisms

**DOI:** 10.1101/2023.09.30.560339

**Authors:** L.L.G. Carrette, A. Santos, M. Brennan, D. Othman, A. Collazo, O. George

## Abstract

Chronic alcohol consumption leads to dependence and withdrawal symptoms upon cessation, contributing to persistent use. However, the brain network mechanisms by which the brain orchestrates alcohol withdrawal and how these networks are affected by pharmacological treatments remain elusive. Recent work revealed that alcohol withdrawal produces a widespread increase in coordinated brain activity and a decrease in modularity of the whole-brain functional network using single-cell whole-brain imaging of immediate early genes. This decreased modularity and functional hyperconnectivity are hypothesized to be novel biomarkers of alcohol withdrawal in alcohol dependence, which could potentially be used to evaluate the efficacy of new medications for alcohol use disorder. However, there is no evidence that current FDA-approved medications or experimental treatments known to reduce alcohol drinking in animal models can normalize the changes in whole-brain functional connectivity. In this report, we tested the effect of R121919, a CRF1 antagonist, and naltrexone, an FDA-approved treatment for alcohol use disorder, on whole-brain functional connectivity using the cellular marker FOS combined with graph theory and advanced network analyses. Results show that both R121919 and naltrexone restored the functional connectivity of the prefrontal cortex during alcohol withdrawal, but through divergent mechanisms. Specifically, R121919 increased FOS activation in the prefrontal cortex, partially restored modularity, and normalized connectivity, particularly in CRF1-rich regions, including the prefrontal, pallidum, and extended amygdala circuits. On the other hand, naltrexone decreased FOS activation throughout the brain, decreased modularity, and increased connectivity overall except for the Mu opioid receptor-rich regions, including the thalamus. These results identify the brain networks underlying the pharmacological effects of R121919 and naltrexone and demonstrate that these drugs restored different aspects of functional connectivity of the prefrontal cortex, pallidum, amygdala, and thalamus during alcohol withdrawal. Notably, these effects were particularly prominent in CRF1-and Mu opioid receptors-rich regions highlighting the potential of whole-brain functional connectivity using FOS as a tool for identifying neuronal network mechanisms underlying the pharmacological effects of existing and new medications for alcohol use disorder.

## INTRODUCTION

Chronic alcohol consumption leads to dependence and withdrawal symptoms upon cessation, contributing to persistent use. Complex behaviors, including those that emerge during alcohol withdrawal, are modulated by various factors, such as sensory input, internal state, motivation, memory, and resulting actions that are coordinated throughout brain-wide changes in activity (1-3). However, the orchestration of alcohol withdrawal by the brain, as a whole, and the effects of pharmacological treatments on these brain networks remained elusive. It is critical to identify the brain network changes caused by alcohol dependence and neuropharmacological interventions to reduce alcohol drinking and relapse in animal models to understand the consequences of alcohol use and identify the mechanisms responsible for the therapeutic efficacy of existing and novel medications. Site-, cell-, and circuit-specific manipulations using lesions, pharmacology, optogenetics, and chemogenetics have identified critical brain regions and circuits responsible for excessive alcohol drinking and the emergence of negative emotional states during withdrawal, including the extended amygdala, striatum, prefrontal cortex, among others (4-25). However, the coordination of brain-wide activity between and within these brain regions during alcohol withdrawal remains poorly understood. Visualizing functional changes in whole-brain networks produced by alcohol dependence, withdrawal, and neuropharmacological interventions to reduce alcohol drinking and relapse is a critical step to identifying the neurobiological mechanisms responsible for the pharmacological effects of these treatments.Leveraging advances in whole-brain immunolabeling, brain clearing, light-sheet microscopy imaging (26), automated detection (27), and graph theory analysis (28) provide an opportunity to bridge this gap and study the functional connectomes of dependent animals in withdrawal from alcohol (1, 2) and stimulants (29, 30). For instance, withdrawal from alcohol, nicotine, cocaine, and methamphetamine leads to coordinated hyperactivity of the extended amygdala network, with a shift from a cortical-driven to a subcortical-driven network, and a reduction of network modularity as a result. Moreover, after resumed access to alcohol, network modularity normalizes relative to abstinence (2). We hypothesized that these network changes may represent relevant biomarkers of withdrawal in dependency and that pharmacological treatments known to show some level of efficacy in animal models relevant to alcohol use disorder should normalize whole-brain functional connectivity. To this end, we examined the effects of two treatments: 1) naltrexone, an FDA-approved mu-opioid receptor (MOR) antagonist, and 2) R121919, an experimental corticotropin-releasing factor receptor 1 (CRF1) antagonist, using single-cell whole-brain imaging, on the whole-brain reactivity and functional connectome of non-dependent and dependent animals during alcohol withdrawal, which revealed specific and opposite effects of the treatments on the dependent brain.

## MATERIALS AND METHODS

### Animals

Male and female C57BL/6J mice (Jackson Laboratories) (N = 48), 8-week-old, group-housed on a 12 h/12 h normal light/dark cycle in a temperature (20-22°C) and humidity (45-55%) controlled vivarium with ad libitum access to tap water and food pellets (PJ Noyes Company, Lancaster, NH, USA). All procedures were conducted in strict adherence to the National Institutes of Health Guide for the Care and Use of Laboratory Animals and were approved by the Institutional Animal Care and Use Committee of UC San Diego.

### Dependence induction

Alcohol dependence was induced by 9 days of daily intraperitoneal (i.p.) injection of EtOH (2 g/kg, 20% w/v in 0.9% saline, or saline control) with the alcohol dehydrogenase inhibitor fomepizole (4-methylpyrazole, 9 mg/kg, Sigma-Aldrich)(31). Blood alcohol level was assessed from tail bleeds 1h (intoxication) and 24h (withdrawal) after injection on day 6 or just before injection on day 7 using gas chromatography. One female from the naltrexone treatment group was euthanized following adverse effects from alcohol.

### Drugs

Naltrexone hydrochloride (Sigma-Aldrich) was dissolved in physiological (0.9%) saline for i.p. injection of 3 mg/kg (0.3 mg/ml) (32-34). R121919 (Tocris) was dissolved in 10% DMSO, 5% cremophor, and 85% water for i.p. injection of 20 mg/kg (2 mg/ml) (35-37). Saline (0.9%) was used as control.

### Withdrawal measurement by digging and marble burying

Digging and marble burying were measured to quantify withdrawal 24 hours after the last EtOH injection and 30 min after drug treatment as described before (1, 38, 39). Testing was conducted under dim lighting (11-20 lux). The mouse was placed in a new, clean cage with a bedding thickness of 5 cm without lid and was allowed to freely dig for 3 min. The number of digging bouts and total digging duration were recorded (phase 1). The mouse was next placed in a similar new cage with 12 marbles evenly spaced on top of the bedding and allowed to bury the marbles for 30 min with a lid. The number of marbles that were covered two-thirds or more by bedding was counted at the end of the test (phase 2) The observers were blinded to the treatment groups.

### Pericardial Perfusion

Brains were collected 90 min after injection. The mice were deeply anesthetized with isoflurane and perfused with ice-cold 15 mL of phosphate-buffered saline (PBS) followed by 30 mL of 4% paraformaldehyde in PBS. The brains were postfixed in 4% paraformaldehyde overnight at 4°C with agitation. The next day, brains were washed 3 times for 30 min with PBS (with 0.1% sodium azide) at room temperature (RT) and stored in the same solution at 4°C until further processing.

### Brain FOS immunolabeling and clearing using iDISCO+ protocol

with small alterations from earlier descripsion (1, 27, 29, 40). Fixed samples were washed in 20% methanol (in milliQ water) for 1 h, 40%methanol for 1 h, 60% methanol for 1 h, 80% methanol for 1 h, and 100% methanol for 1 h twice. The samples were then precleared with overnight incubation in 33% methanol/66% dichloromethane (DCM). The next day, the samples were washed 2x in 100% methanol for 3 h and then bleached with 5% H2O2 in methanol at 4°C overnight. After bleaching, the samples were rehydrated in 80% methanol (in milliQ water) for 1 h, 60% methanol for 1 h, 40% methanol for 1 h, 20% methanol for 1 h, PBS for 1 h, and PBS/0.2% TritonX-100 for 1 h twice. The samples were then permeabilized by incubation in PBS/0.2% TritonX-100/20% dimethyl sulfoxide (DMSO)/0.3 M glycine at 37 °C for 2 days (d) and blocked in PBS/0.2% TritonX-100/10% DMSO/6% donkey serum at 37 °C for 2 d. The samples were next incubated in rabbit anti-c-FOS (1:2000, 226 008; Synaptic Signaling) in PBS/0.2% Tween with 10 μg/mL heparin (PTwH)/5% DMSO/3% donkey serum at 37°C for 7 d.The samples were then washed in PTwH for 24 h (5x) and incubated in donkey anti-rabbit Alexa647 (1:500, A31573; Invitrogen) in PTwH/3% donkey serum at 37°C for 7 d. Finally, the samples were washed in PTwH for 24 h (5x) before clearing and imaging, for the samples were dehydrated in 20% methanol (in milliQ water) for 1 h, 40% methanol for 1 h, 60% methanol for 1 h, 80% methanol for 1 h, 100% methanol for 1 h, and 100% methanol again overnight. The next day, the samples were incubated for 3 h in 33% methanol/66% DCM. The methanol was then washed out twice in 100% DCM for 1 h, or until the brains sank in the solution. Finally, the samples were incubated in dibenzyl ether (DBE; 108014-1KG; Sigma) for minimum 4 h and then stored in fresh DBE at RT in the dark until imaged.

### Light-sheet microscopy imaging

Left hemispheres of cleared samples were imaged in the sagittal orientation (left lateral side up) on a light-sheet microscope (Ultramicroscope II; LaVision Biotec) equipped with a scientific complementary metal–oxide–semiconductor camera (Andor Neo), 2×/0.5 objective lens (MVPLAPO 2×), and 6-mm working distance dipping cap. Imspector microscope controller v144 software was used. The microscope was equipped with an NKT Photonics SuperK EXTREME EXW-12 white light laser with three fixed light-sheet-generating lenses on each side. Scans were made at 0.8× magnification (1.6× effective magnification) with a light-sheet numerical aperture of 0.148. Excitation filters of 480/30, 560/40, and 630/30 were used. Emission filters of 525/50, 595/40, and 680/30 were used. The 560-nm channel was just used to confirm the labeling in the 630-nm channel was specific. All samples were scanned with a Z-step size of 3 μm using two-sided illumination (combined into one image using the blending algorithm). For the 630-nm channel, dynamic horizontal scanning was used (10 to 20 acquisitions per side with exposures ranging from 150 to 800 ms, combined into one image using the horizontal adaptive algorithm). For the 480-nm channel, no horizontal scanning was used with exposures ranging from 90 to 200 ms. To accelerate acquisition, both channels were acquired in two separate scans. To account for micromovements of the samples that may occur between scans, 3D image affine registration was performed to align both channels using ClearMap (27).

### Image processing

Light-sheet microscope images were analyzed from the end of the olfactory bulbs (the olfactory bulbs were not included in the analysis) to the beginning of the hindbrain and cerebellum. Counts of FOS-positive nuclei from each sample were identified for each brain region using ClearMap (27). ClearMap uses autofluorescence that is acquired in the 488 channel to align the brain to the Allen Mouse Brain Atlas (41) and then registers FOS counts to regions that are annotated by the atlas. The data were log10 normalized to reduce variability. One brain from the dependent naltrexone group was excluded due to a problem with alignment.

### Statistical analysis

The significance level was set for p at 0.05. Calculations and visualization were performed in Prism Graphpad and R. Behavioral data and network centrality measures were analyzed using 2-way ANOVA for treatment and alcohol dependency. Total FOS counts were analyzed between groups with one-way ANOVA followed by Dunnett’s post-hoc test for discreet data compared to a control. The test was performed comparing all groups to the control saline treated group and comparing the dependent treated groups to the dependent saline treated groups. For comparison of FOS counts between groups and regions, the same test was performed but the p values were corrected using the Benjamini-Hochberg principle for false discovery rate with q at 10%. Percent-change in FOS counts between groups and regions were also compared by calculating the 95% confidence interval through bootstrap resampling with replacement stratified per group using the ‘boot’ package (42). The “boot’ package was also used to estimate the standard error of the mean for the network modularity, with N=100 for resampling. LASSO multiple regression analysis was performed using the ‘glmnet’ package (43), with alpha=1 and optimizing lambda using tuneGrid. Plots were made with the ‘ggplot’ package (44), heatmaps were made with the ‘complexHeatmaps’ package (45). Analysis of average correlation of distinct brain regions with the rest of the brain was performed by 2-way ANOVA between treatments and regions with Bonferroni corrected post-hoc tests.

### Network analysis

Pearson correlations of the normalized counts between all brain regions (nodes) were calculated within the treatment groups. Brain regions could then be hierarchically clustered (method = complete) using the complete Euclidean distance based on the correlation values (edges). The clustering dendrograms were trimmed at half the height of each given tree to split the dendrogram into specific modules. The network could also be thresholded to remove any edges that were weaker than R = 0.75. The centrality measures degree and betweenness were calculated using a customized version of the bctpy Python package (https://github.com/aestrivex/bctpy), which is derived from the MATLAB implementation of Brain Connectivity Toolbox (46). Minimal networks were obtained from the nodes and edge tables using Gephi software (47).

### ISH expression analysis

Structure and gene expression data were extracted from the In Situ Hybridization gene expression database and Allen Brain Atlas (48) in Python, as published and described before (30, 49-51), and intersected with the brain regions from the FOS dataset.

## RESULTS

### Animal model relevant to alcohol dependence and single-cell whole-brain imaging pipeline

To test the effect of pharmacological treatments on alcohol withdrawal, mice first received 9 intraperitoneal (2 g/kg, *i.p*.) alcohol injections to induce dependence (31). Each injection raised average blood alcohol levels >200 mg/dL in 1 h, which returned to 0 mg/dL within 24 h (**Fig. 1A-C**). On day 10, 24 h after the last injection, mice received R121919 (20 mg/kg *i.p*.), naltrexone (3 mg/kg *i.p*.), or saline and underwent assessment for withdrawal-related behaviors using digging and marble-burying paradigms (38, 39). The dependent animals exhibited significantly more withdrawal-induced bouts of digging (2-way ANOVA, F(1,41)=5.7, p=0.022, **Fig. 1D**) and longer digging durations (2-way ANOVA, F(1,41)=9.5, p=0.004, **Fig. 1E**), but buried a similar number of marbles (2-way ANOVA, F(1,41)=1.6, p=0.2, **Fig. 1F**) compared to control animals. The treatments did not cause significant changes in the behavior within the dependent or control animals (2-way ANOVA, F(2,41)<1, p>0.4 for treatment; 2-way ANOVA, F(2,41)<0.2, p>0.8 for interaction). These results show that following the chronic alcohol treatment the mice exhibited alcohol withdrawal-induced digging.

**Figure 1.**
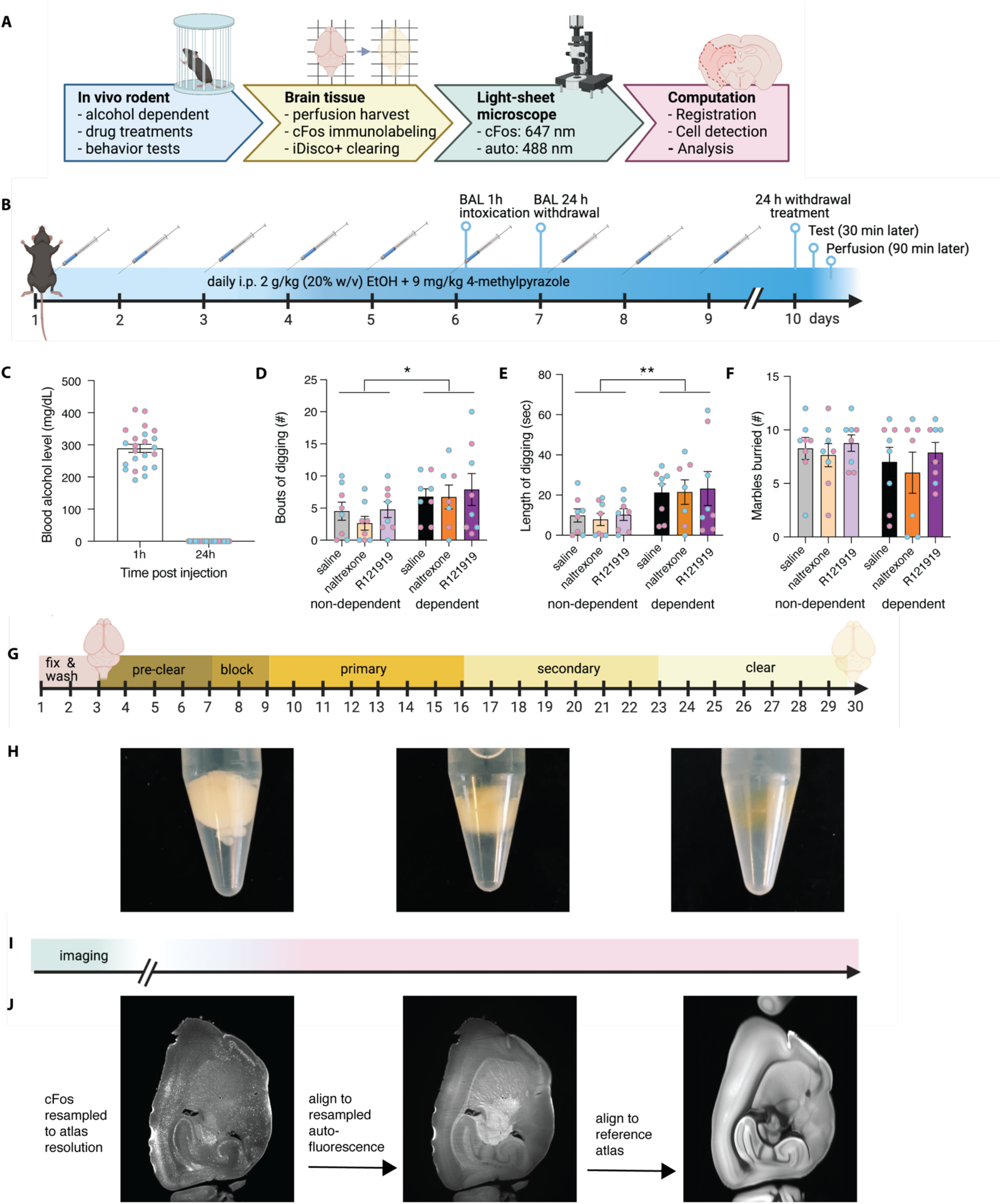
Experimental overview. **(A)** General timeline of the experiments. **(B)** Detailed timeline of the in vivo experiments: 9 daily alcohol injections, collections for blood alcohol levela (BAL) at intoxication and withdrawal, 1h and 24 h following injection, treatments on day 10, 24 h in withdrawal, with digging tests after 30 min and perfusion 90 min later. **(C)** BAL results for intoxication (1 h) and withdrawal (24 h). **(D-F)** Behavior data of animals following treatment with R121919 (purple), naltrexone (orange) or saline (black) treatment following chronic alcohol (or saline control, in lighter hues) for bouts of digging (D), length of digging (E), and number of marbles buried (F) (males = blue, females = pink). Analyzed by 2-way ANOVA, *p<0.05, **p<0.01. **(G)** Detailed timeline of brain immunolabeling and clearing. **(H)** Representative images of the brains in the last step of the protocol turning transparent through refraction index matching in dibenzylether. **(I)** Timeline of imaging and image processing with resampling and aligning FOS to autofluorescene to reference atlas.

To visualize the alcohol withdrawal brain state and the effect of the treatments in these mice, 90 minutes after the treatment, brains were perfusion-fixed, harvested, and subjected to the iDisco+ pipeline for FOS immunolabeling and clearing (**Fig. 1G**). The last step rendered the brains transparent (**Fig. 1H**) allowing for full imaging at single-cell resolution with light-sheet microscopy (**Fig. 1H**). The automated ClearMap pipeline can automatically detect, count, and register all FOS-positive cells in the regions of the Allen Brain Atlas (**Fig. 1J**) (27), which results in lists of FOS-positive cell counts per brain region for each brain. Using LASSO multivariate linear regression modeling, the brain regions that contributed most to each behavior based on the individual regional FOS reactivity can then be identified (43) (See **Fig. S1**). In short, the whole-brain reactivity, as represented by FOS-positive cells was counted per region.

### Alcohol withdrawal increases while naltrexone reduces whole-brain neuronal reactivity

To understand differences in neuronal reactivity following both chronic alcohol treatment and single dose treatment with R121919 and naltrexone in alcohol-dependent and non-dependent controls, FOS counts were compared (**Fig. 2A**). Looking at the total whole-brain FOS counts, both treatment with either R121919, naltrexone, or alcohol caused an upward trend compared to non-dependent, saline treatment (one-way ANOVA, F(5,40)= 2.57, p=0.04), but Dunnett post-hoc did not reveal any significant differences). Once alcohol-dependent, naltrexone treatment significantly reduced the total FOS count compared to dependent saline-treated controls (one-way ANOVA, F(2,19)= 4.70, p=0.02, with Dunnett post-hoc p=0.05). R121919 did not cause a significant change in Fos counts. These results reveal a dependent brain state with increased total reactivity that is normalized with naltrexone.

**Figure 2.**
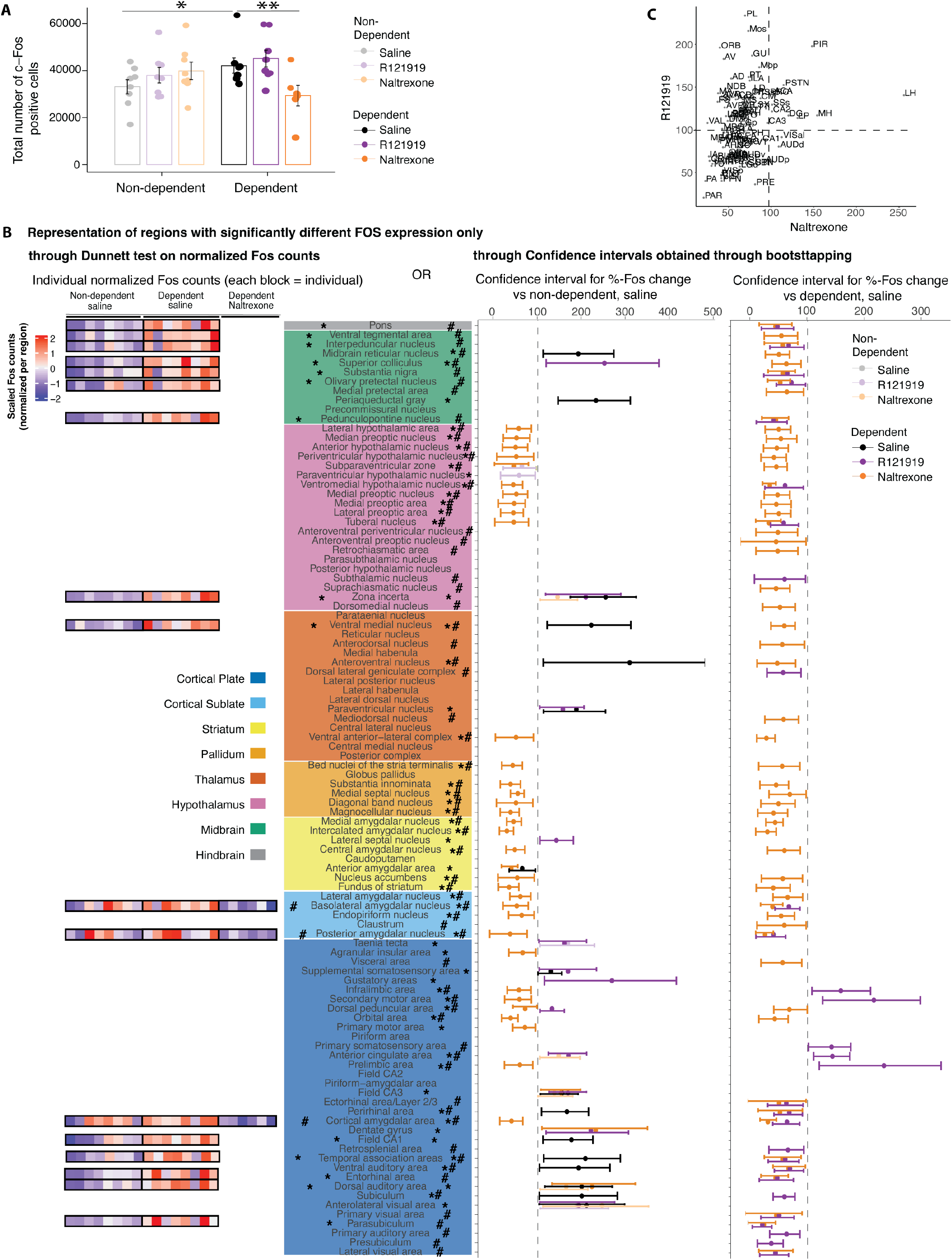
*Brain-wide reactivity through FOS counts and %-change:* Alcohol withdrawal increases, naltrexone reduces whole-brain neuronal reactivity. **(A)** Total FOS counts between groups (*p=0.04, **p = 0.02 with one-way anova). **(B)** Regional FOS counts that are significantly different through 2 different approaches: q<0.1 False Discovery Rate on Dunnett test (*saline-treated alcohol dependent vs. non-dependent, #Nalterxone-treated dependent vs. saline-treated dependent; left). Bootstrapped 95-% confidence intervals (for *relative FOS %-change of naltrexone, R121919, and alcohol treatments vs. saline-treated control animals; middle or #naltrexone and R121919 treatments in dependent animals vs. saline-treated dependent animals; right). **(C)** Relative FOS %-change of naltrexone (x) and R121919 (y) treatments in dependent animals versus saline-treated dependent animals.

To understand the differences in FOS reactivity between the main brain regions (n=100), differences in FOS counts per region were next compared (**Fig. 2B** and **S2**). One-way ANOVA with Dunnett post-hoc performed on the FOS counts between the different groups as above, but with additional Benjamini-Hochberg false discovery rate adjustment (q=10%) to correct for the multiple comparisons, identified 14 regions that were significantly activated in dependent animals compared to controls (**Fig. 2B** left* and showing individual FOS counts normalized per region and **Fig. S3**, top), mainly in the midbrain, with the ventral tegmental area (VTA), interpeduncular nucleus (IPN), superior colliculus (SC), substantia nigra (SN), olivary pretectal nucleus (OP), pedunculopontine nucleus (PPN) and cortical plate with the field CA1 (CA1), temporal association areas (TEa), entorhinal area (ENT), dorsal auditory area (AUDd) and parasubiculum (PAR), in addition to the pons (P) in the hindbrain, zona incerta (ZI) in the hypothalamus, and ventral medial nucleus (VM) of the thalamus. Three amygdalar regions, the basolateral amygdalar nucleus (BLA), the posterior amygdalar (PA) nucleus and the cortical amygdalar nucleus (COA), were found to be significantly (q<0.1) downregulated following naltrexone treatment in dependent animals compared to saline treatment (**Fig. 2B** left# and showing individual normalized FOS counts and **Fig. S3**, bottom). These regions had a trend to be upregulated from control animals following alcohol treatment, to then be significantly reduced with naltrexone treatment.

A second way to add statistical insight to the data is to determine bootstrapped confidence intervals of the populations. Repeat sampling with replacements from the test and control samples (saline-treated dependent or non-dependent), allows for the construction of 95%-confidence intervals for the %-change of FOS counts. Confidence intervals that do not include 100%, or in other words no change, are considered significantly different (**Fig. 2B** middle, representation of significantly different intervals only, all intervals, including non-significant ones can be found in **Fig S4**). This analysis confirmed the upregulation of FOS reactivity in dependent animals during withdrawal compared to controls, through 15 regions mostly overlapping with the ones identified above in the midbrain (midbrain reticular nucleus (MRN), periaqueductal gray (PAG)), and cortical plate (CA1, CA3, TEa, perirhinal area (PERI), subiculum (SUB), AUDd, ventral auditory area (AUDv), anterolateral visual area (VISal), supplemental somatosensory areas (SSs)), in addition to the hypothalamus (ZI) and thalamus (VM and anteroventral nucleus (AV)). There was also 1 region, the anterior amygdalar area (AAA), with down-regulation of the reactivity in alcohol dependent animals compared to non-dependent controls. Naltrexone and R121919 treatments of non-dependent mice caused much fewer changes.Naltrexone increased reactivity in 5 regions in the cortical plate (ACA, CA3, AUDd, and VISal) and thalamus (ZI), while R121919 caused 2 up-regulations in the cortical plate (VISal, and tenia tecta (TT)) and 2 down-regulations in the hypothalamus (subparaventricular zone (SBPV) and paraventricular hypothalamic nucleus (PVH)).

In dependent animals following naltrexone treatment there was a general downregulation of FOS reactivity in 56 brain regions, compared to saline-treated dependent animals (**Fig. 2B** right, representation of significantly different intervals only, all intervals, including non-significant ones can be found in **Fig S4**), again, mostly located in the amygdala (CoA, BLA, medial amygdalar nucleus (MEA), PA, LA), hypothalamus, (lateral hypothalamic area (LHA), medial preoptic area (MPO), MPN, suprachiasmatic nucleus (SCH), ventromedial hypothalamic nucleus (VMH), anterior hypothalamic nucleus (AHN), lateral preoptic area (LPO), periventricular hypothalamic nucleus (PV), subparaventricular zone (SBPV), tuberal nucleus (TU)), pallidum (magnocellular nucleus (MA), medial septal nucleus (MS), diagonal band nucleus (NDB), substantia innominata (SI)), and midbrain (VTA, IPN, OP, SC, SN, medial pretectal area (MPT)), midbrain reticular nucleus (MRN)). R121919 also caused downregulation of FOS reactivity in 24 regions, especially those that saw an upregulation with alcohol intake like the hindbrain P and the temporal areas (Sub, PAR, Ent, TEa, PA, CoA). But contrary to naltrexone, R121919 also caused FOS upregulation in 5 regions including the cortical regions in the prefrontal cortex with the anterior cingulate area (ACA), infralimbic area (ILA), prelimbic area (PL), and the somatosensorimotor cortices with the primary somatosensory area (SSp) and secondary motor area (Mos).Finally, when comparing the treatments in dependent animals to the saline-treated non-dependent rather than dependent saline-treated controls revealed less changes (**Fig. 2B** middle). Naltrexone had 34 downregulated regions and 3 upregulated, R121919 had 12 upregulated regions.

Overall, these analyses showed that alcohol dependence caused widespread changes in regional neuronal reactivity, mostly upregulation, which is reversed by naltrexone treatment. R121919 on the other hand increased reactivity in some and decreased reactivity in other brain regions (**Fig. 2C**).

### Partial normalization of network modularity after treatment with R121919 in alcohol dependent mice

To understand the effect of the treatments on the alcohol withdrawal-induced modifications of whole-brain functional connections between brain regions, the functional connectivity was approximated by correlation in the FOS counts between those brain regions, as described before (1). The modularity of the functional connectomes could then be derived by hierarchical clustering of the regions and cutting the clustering dendrogram at half height (**Fig. 3A-H**, dotted line). The default non-dependent saline network had 12 modules (**Fig. 3A**). Treatment of control animals with R121919 and naltrexone only slightly affected the number of modules (n=9 and 11, respectively, (**Fig. 3B-C**)). The number of modules went significantly down in the network of dependent animals in withdrawal (n=5, **Fig. 3D**, 2-way ANOVA with SEM derived through bootstrapping with N=100, F(1,40)=9.978, p=0.02 for effect of chronic alcohol vs saline). While there was no significant effect of treatments or interaction with alcohol (2-way ANOVA with SEM derived through bootstrapping with N=100, F(2,40)=0.5512, p=0.58 and F(2,40)=2.573, p=0.089 respectively), the effect of treatment in dependent animals had an opposite trending effect on the modularity. Treatment with R121919 seemed to more normalize the network and restore the modularity (n=9, which is the same as non-dependent animals treated with R121919, **Fig. 3E**). Treatment with naltrexone seemed to further reduce the modularity, resulting in a few large modules with strongly correlated brain regions (n=3, **Fig. 3A**). The colored sidebar indicates which anatomical group each brain region belongs to and illustrates thorough reorganization.Independent of the set height definition for cutting off the modules, withdrawal in alcohol-dependent animals caused a reduction in modularity, which was partially normalized after treatment with R121919, but further reduced after treatment with naltrexone (**Fig. 3G-H**).

**Figure 3.**
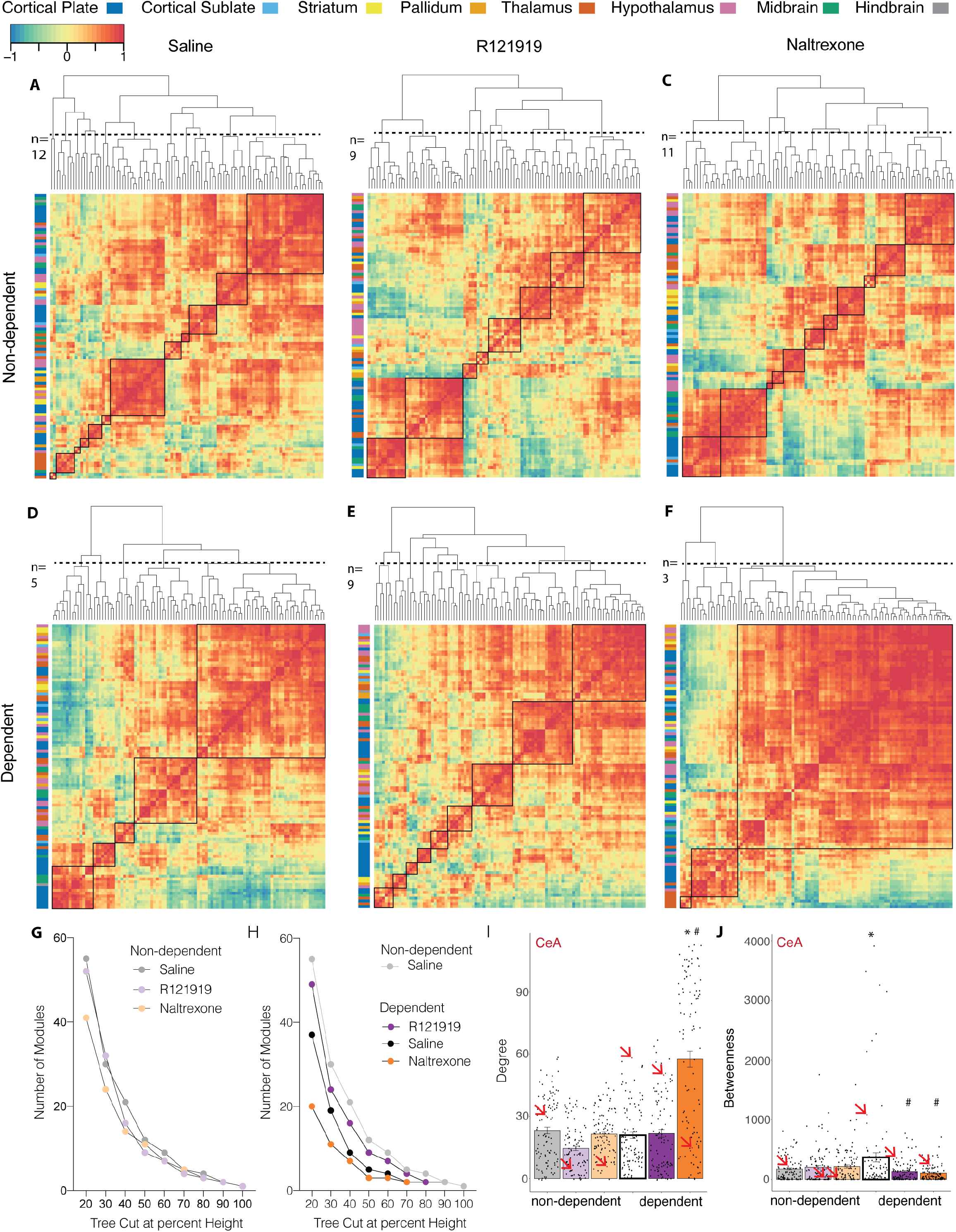
Partial normalization of network modularity after treatment with R121919 in alcohol-dependent mice. **(A-F)** Network modules obtained through hierarchical clustering of regions based on their correlation and cutting the dendrogram at half-height (dotted line, n=number of clusters) in correlation heatmaps from blue (= anticorrelated) to red (= correlated): for control mice treated with saline (A), R121919 (B), or naltrexone (C) and alcohol-dependent mice in withdrawal treated with saline (D), R121919 (E), or naltrexone (F); sidebar indicates the anatomical group the region belongs to: cortical plate (blue), cortical subplate (light blue), striatum (yellow), pallidum (orange), thalamus (red), hypothalamus (pink), midbrain (green), hindbrain (gray). The specific order of the regions in these hierarchically clustered heatmaps can be found in the supplementary information (**Supp. Table 2**). **(G-H)** Number of modules as a function of tree cut for control animals (G) and alcohol-dependent animals (H). **(I-J)** Network centrality measures identifying the CEA (red dot and arrow) as a hub region in the dependent, saline-treated network with high degree (I) and betweenness (J); *p<0.001 compared to non-dependent and #p<0.001 compared to dependent saline-treated animals).

To understand other network properties of the functional connectome centrality measures were calculated: the degree or the number of connections of a region (**Fig. 3I**) and the betweenness or the number of shortest paths between any two regions that go through a region (**Fig. 3J**). The average and maximum degree in naltrexone-treated dependent animals was higher than both the non-dependent and dependent saline controls (Bonferroni corrected post-hoc p<0.001 following a significant 2-way ANOVA interaction between treatment and dependence F(2,594)=47.89 with p<0.001) due to strongly increased correlations. In contrast, following R121919 treatment of dependent animals, the degree remains very similar to the saline-treated animals (Bonferroni corrected post-hoc p>0.99). For betweenness, naltrexone and R121919 both reduced the average network betweenness compared to saline-treated dependent animals (Bonferroni corrected post-hoc p<0.001 following a significant 2-way ANOVA interaction between treatment and dependence F(2,594)=10.38 with p<0.001), which was elevated compared to saline-treated controls (Bonferroni corrected post-hoc p<0.01). Treatments had no effects in non-dependent animals. Brain regions that have both a high degree and high betweenness can be considered hub regions that play central roles in the network (28). The CEA fulfilled these criteria as a network hub of the saline-treated dependent animals, but was no longer within the top 10 of regions for degree and betweenness following R121919 treatment, and was not even amongst the top regions anymore following naltrexone treatment.

F (2, 40) = 0.5512

These results illustrate, that both R121919 and naltrexone strongly affected the network properties of the functional connectome in alcohol-dependent animals in withdrawal, which was significantly altered from non-dependent control animals, but with an almost opposite effect.

### Receptor-expressing regions drive the treatment-induced network changes in dependent mice

For easier comparison of the whole brain effects of alcohol withdrawal, treatment with R121919 and naltrexone, the networks were reorganized to list the brain regions in the same order according to the major anatomical groups (**Fig. 4A**; exact order in **table S3**). We further aligned the FOS expression of all the regions relative to non-dependent saline-injected for dependent saline-injected animals and relative to dependent saline-injected for the dependent R121919 and naltrexone treated. Finally, we also aligned the whole-brain expression profile of the receptors R121919 and naltrexone act on, corticotropin-releasing hormone receptor 1 (CRF1) and mu opioid receptor respectively, as extracted from the Allen Brain Atlas ISH Database (48). The reduced modularity of the saline-treated dependent animals resulted from an overall increased reactivity and functional connectivity, including hyperconnectivity to the frontal cortex, but hypoconnectivity to the sensory cortex. (**Fig. 4B**) Following R121919 treatment, the distinctive connectivity in the cortical plate was normalized (**Fig. 4C**). This was the result of earlier shown diminished FOS expression and reactivity in the sensory cortex, including TEa and PAR, where it was elevated by alcohol withdrawal, and the increased FOS expression and reactivity in the frontal cortex, including the ILA and PL. Note that these regions driving the normalization had the highest expression levels of *Crhr1*, the gene coding for CRF1, which was mostly expressed in the cortical plate; top levels include PL, TEa, auditory and visual cortex. Following naltrexone treatment, most brain regions reduced FOS expression and reactivity, resulting in increased functional connectivity (**Fig. 4D**). Except for brain regions in the thalamus that became more disconnected, including the habenula. Note that these regions had amongst the highest expression levels of *Oprm1*, the gene coding for MOR, which was mostly expressed in the thalamus; top levels include MH, LH, PVT and central medial nucleus of the thalamus. Naltrexone also works on the kappa opioid receptor (KOR), which is mainly expressed in the cortical plate and subplate, which could explain the expansion of the network to include the sensory cortex (**Fig. S4**). These results illustrated that the brain regions expressing most of the targeted receptors, have the biggest reactivity differences and drive the treatment-induced network changes.

**Figure 4.**
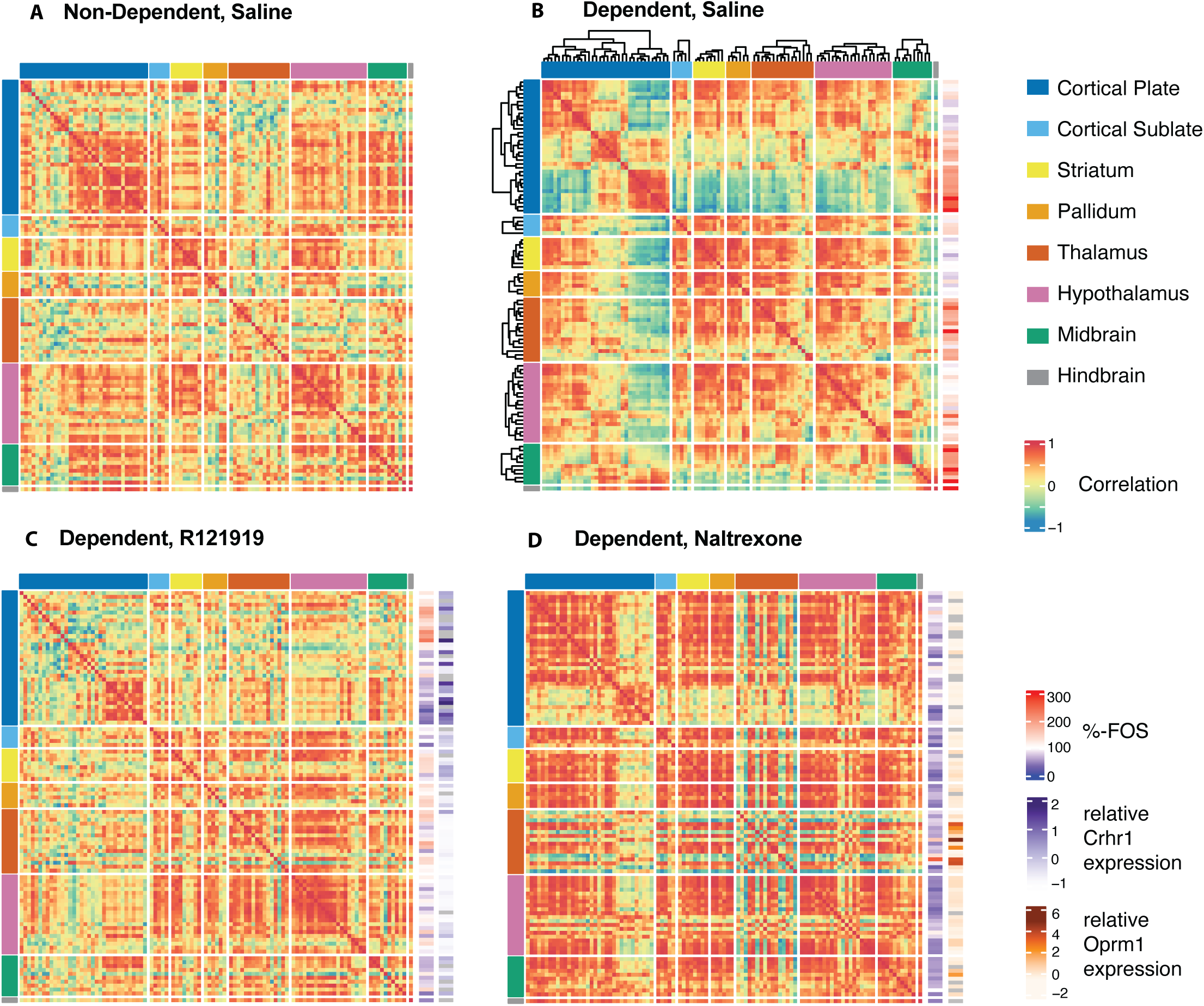
Whole-brain Crhr1 and Oprm1 mRNA expression aligned with FOS expression changes and functional connectomes as correlation heatmaps from blue (= anticorrelated) to red (= correlated) with regions hierarchically organized within the major anatomical subgroups for saline-treated dependent animals: in non-dependent saline-treated animals (A), alcohol-dependent animals saline-treated, with relative FOS expression to non-dependent saline-treated (B), R121919-treated with FOS expression to dependent saline-treated and Crhr1 expression (purple) (C), or naltrexone-treated with FOS expression to dependent saline-treated and Oprm1 expression (orange) (D). The order of the brain regions for this figure can be found in supplemental table 3.

### R1219191 excludes from, while naltrexone integrates the prefrontal cortex in the addiction network

To approach the data in a biased fashion, regions known to be relevant to alcohol dependence can be focused on to reveal the connections between them complementing the non-biased approach. Regions of known importance to addiction include 1) the amygdala, the central nucleus of the amygdala was found as the central hub of the alcohol-dependent withdrawal network (**Fig. 3i)**; here we looked at the posterior amygdalar nucleus (PA), basolateral amygdalar nucleus (BLA), cortical amygdalar area (CoA), central amygdalar nucleus (CEA), medial amygdalar nucleus (MEA), anterior and amygdalar area (AAA)(**Fig. 5**, labeled red); 2) the regions of the dopamine-regulated positive reinforcing circuitry, consisting of the nucleus accumbens (ACB), caudoputamen (CP), lateral hypothalamic area (LHA), and ventral tegmental area (VTA) (**Fig. 5**, labeled green); 3) the regions of the stress-related negative reinforcing circuitry, consisting of the bed nuclei of the stria terminalis (BST), medial habenula (MH), lateral habenula (LH), paraventricular hypothalamic nucleus (PVH), and interpeduncular nucleus (IPN) (**Fig. 5**, labeled blue); and finally 4) the regions controlling executive functions in the frontal cortex, including the prelimbic area (PL), infralimbic area (ILA), anterior cingulate area (ACA), dorsal peduncular area (DP), agranular insular area (AI), and orbital area (ORB) (**Fig. 5**, labeled in turquoise).Focusing on the miniature connectome amongst these regions revealed shifts in communication (**Fig. 5A-D**). While the addiction network in the non-dependent brains was overall well-connected, with the exception of the habenula (MH and LH; **Fig. 5A**), some separation occurred in dependent animals. The connection of some clusters was reduced from the rest of the network, like the frontal cortex and some parts of the amygdala, but the MH became more integrated (**Fig. 5B**). The frontal cortex further disconnected and the habenula further integrated following R121919 treatment (**Fig. 5C**). Following treatment with naltrexone, on the other hand, the opposite occurred. The habenula and PVH, regions known to contribute to the negative emotional states became completely disconnected, while the frontal cortex was strongly integrated (**Fig. 5D**).

**Figure 5.**
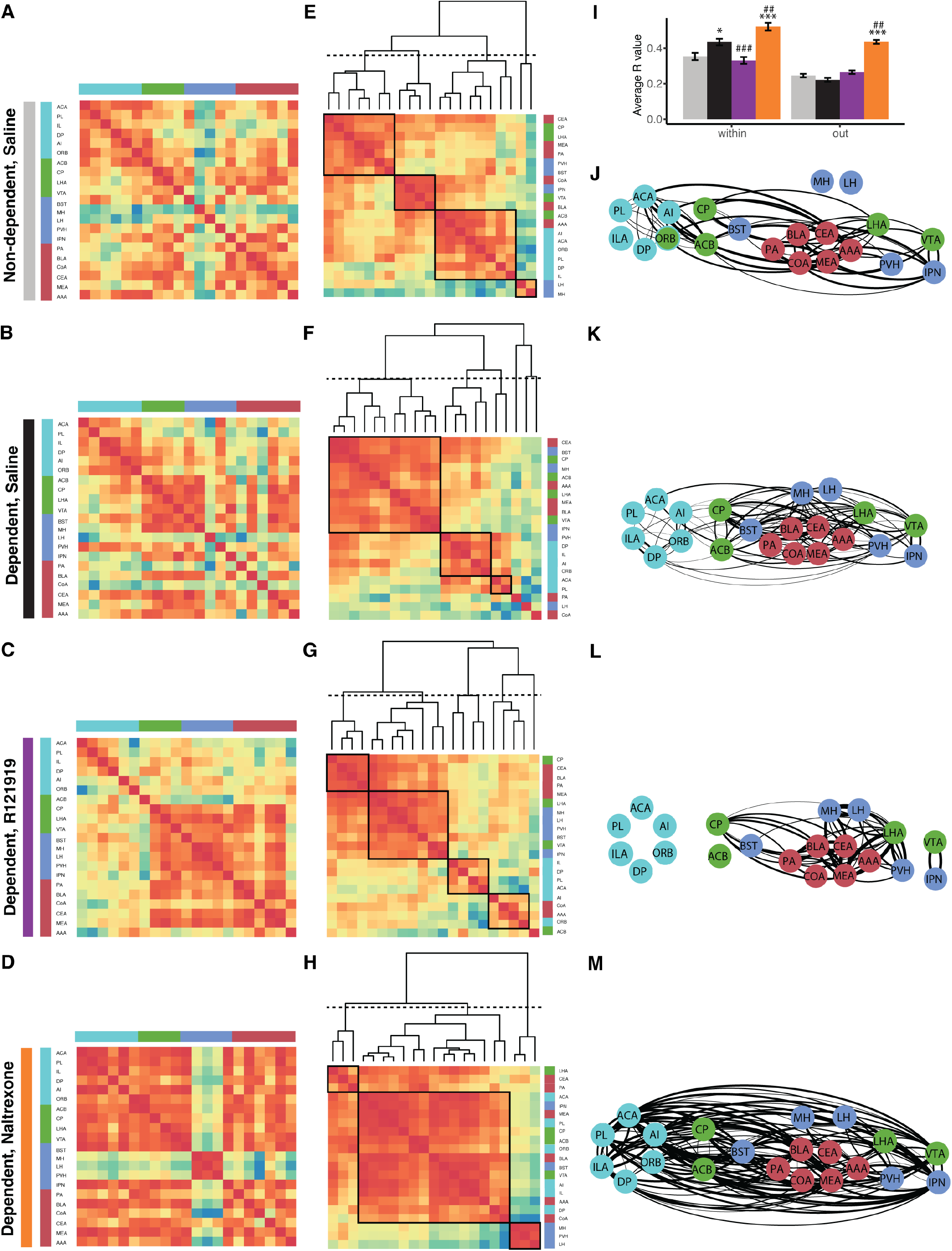
The dependency-induced changes in the selected minimal addiction network are targeted by the treatments. (**A-D**) Selected correlation heatmap, organized through regions of the frontal cortex (turquoise), associated with positive (green) and negative (blue) emotional state, and the amygdala (red); (**E-H**) Reorganized correlation heatmaps by hierarchical clustering; (**I**) Average correlation within the addiction network and from the addiction network with the rest of the brain with *p = 0.02 and ***p<0.001 compared to non-dependent and p = 0.001 and 0.01, respectively, compared to saline-treated dependent with Bonferroni-corrected posthoc ##p = 0.01 and ###p =0.001, compared to saline-treated dependent. (**J-M**) Network representation with the functional distance calculated based on the correlation (R>0.75 represented); for non-dependent saline-treated ((A, E, I), dependent saline-treated (B, F, J), R121919 treated (C, G, K), and naltrexone treated (D, H, L) animals.

The nature of how the miniature connectome changed under the different treatments could further be illustrated by hierarchical clustering of these mini networks (**Fig. 5E-H**) and using network representations with the regions as nodes and the correlation (R>0.75) as the edges between them (**Fig. 5J-M**, for ease of identifying and comparing regions, the location of the nodes in the network was fixed; for the networks with functional positioning of the nodes following the application of a force field, see **Fig. S5**). On average the connectivity within the addiction networks was significantly different between treatments (1-way ANOVA, F(3,1760)=19.36 with p<0.0001) with chronic alcohol treatment significantly increasing the connectivity (p = 0.02 compared to non-dependent with Bonferroni-corrected posthoc) (**Fig. 5I**). R121919 treatment significantly reduced this connectivity and naltrexone treatment significantly increased it further (p = 0.001 and 0.01, respectively, compared to saline-treated dependent with Bonferroni-corrected posthoc). The average connectivity of the addiction network with the rest of the brain was significantly lower than within (2-way ANOVA, F(1,8392)=108, p<0.0001) with an interaction with the treatment (F(3,8392)=8.39, p<0.0001). The overall connection of the addiction network with the rest of the brain does not significantly change in dependent animals compared to non-dependent. However, when looking in detail there is a significant connectivity reduction to the cortical plate and an increase to the thalamus, driven by the amygdala and positive reinforcing circuitry (**Fig. S6-7**). There is also no significant overall change following R121919 treatment in dependent animals compared to saline treatment (**Fig. 5I**). Looking in more detail, there is a significant connectivity increase to the cortical plate and decrease to the striatum, driven by the amygdalar and negative reinforcing circuitry or prefrontal cortex and negative reinforcing circuitry respectively (**Fig. S6-7**). The overall connectivity following naltrexone treatment is significantly increased compared to both non-dependent and dependent animals ((**Fig. 5I**), 1-way ANOVA, F(3,6632)=86.93 with p<0.0001, p<0.0001 with Bonferroni-corrected posthoc). This increase is especially pronounced to the cortical plate, hypothalamus and midbrain and driven by all but the negative reinforcing circuitry (**Fig. S6-7**).

This showed that, on the one hand, the disconnect of the frontal cortex with the rest of the addiction network that was introduced in dependent networks, was completely cut-off following R1219191 treatment and patched following naltrexone treatment. Moreover, the integration of the habenula with the rest of the addiction network that was introduced in dependent networks, was even more integrated following R1219191 treatment and fully disconnected following naltrexone treatment. On the other hand, the increased average functional connectivity within the addiction network in dependent animals following chronic alcohol administration compared to non-dependent controls, was normalized by R121919 treatment, while it was exaggerated by naltrexone treatment.

## DISCUSSION

Here, we tested the effect of a single dose of R121919, a CRF1 antagonist, and naltrexone, an FDA-approved treatment for alcohol use disorder, on whole-brain functional connectivity of alcohol-dependent and non-dependent animals. Dependent animals exhibited withdrawal-induced digging behavior, that was not affected by the treatments. Using single-cell whole-brain imaging, we discerned the effects of the treatments on the alcohol withdrawal brain state in dependent animals and non-dependent controls. Overall, alcohol withdrawal increased brain reactivity, resulting in a widespread network coactivation pattern with reduced network modularity in dependent compared to non-dependent controls. Treatment with R121919 partially normalized the withdrawal-induced network hypomodularity, by acting on CRHR1-rich cortical regions. This resulted in reduced reactivity in the sensory cortex, which was overactive due to withdrawal, and increased reactivity in the prefrontal cortex. Specifically, within the addiction network, R121919 normalized the connectivity that had been enhanced by withdrawal through further disconnection of the prefrontal cortex, but integration of the habenula. In contrast, naltrexone treatment further decreased network modularity in dependent animals through a significant global downregulation in activity. This led to increased correlations between all regions, except the MOR-rich thalamus. Specifically, within the addiction network, naltrexone resulted in hyperconnectivity, with the prefrontal cortex included, but the habenula excluded.

Immediate early genes, such as c-Fos, are rapidly transcribed in response to cellular stimuli, enabling the visualizing of reactivity (52). Furthermore, the principle of “neurons that wire together, fire together” allows us to derive the functional connectome from this reactivity (27, 28). The synchronized neural firing or co-reactivity patterns define a brain state. Since immediate early gene expression integrates neuronal activation over a 1-2 h period, it is ideally suited for identifying brain networks associated with prolonged states like alcohol withdrawal. Consistent with prior research (2, 3), chronic alcohol and withdrawal induced reactivity in several regions of the cortical plate, hypothalamus, thalamus, and especially the midbrain, while reducing functional network modularity (1, 53). Complex networks, like the brain, exhibit a modular organization that optimizes adaptability, efficiency, and robustness while minimizing costs (54). Decreased modularity may be partially responsible for the cognitive dysfunction that is seen in humans and animal models of alcohol dependence (55-57).

In dependent animals, naltrexone downregulated reactivity throughout the brain, particularly in the amygdala and hypothalamus; R121919 downregulated reactivity in the cortical sensory and temporal areas that had become upregulated during withdrawal and upregulated reactivity in the prefrontal cortex compared to saline-treated dependent animals. Comparing the treatments in the dependent animals to the saline-treated non-dependent animals revealed fewer changes, presumably because the treatments restored the dependent brain towards a more baseline state. Additionally, changes were predominantly specific to the dependent state, as treatments in non-dependent animals resulted in far fewer changes. These treatments thus target the dysregulated reactivity and network of the dependent animals. Notable regions downregulated by naltrexone included LA and MPN, which were identified as major contributors to digging bouts and duration through multiple linear regression modeling (**Fig. S1)**, and the COA was recently identified to normalize its reactivity with re-access to drinking in dependent animals and when chemogenetically silenced reduce drinking in withdrawal (2).

The alcohol-dependent network hypomodularity was partially normalized following treatment with R121919. Naltrexone, on the other hand, further reduced modularity, seemingly exacerbating the withdrawal state. However, other network centrality measures potentially hinted towards another conclusion. While the withdrawal network in dependent animals had an increased average betweenness with a similar degree compared to the non-dependent network, naltrexone treatment in dependent animals was associated with a decreased average betweenness and increased degree. High betweenness with a standard degree indicates a strongly organized network with a limited number of orchestrating central hubs. The CEA plays this central role during alcohol withdrawal (4-16) and was found as a hub of the alcohol-dependent network. Conversely, the naltrexone network had strongly increased connectivity, resulting in decreased betweenness and loss of orchestration through central hubs. The contribution of the CEA was reduced following R121919 and completely lost following naltrexone treatment in dependent animals. The role of the CEA was also reduced following R121919 and naltrexone treatment in control animals.

The two treatments evoked nearly opposite effects on the withdrawal network across all measures, which aligns with their function targeting different receptors that play distinct roles in ethanol reinforcement (58). R121919, a CRHR1 antagonist, has previously demonstrated efficacy in reducing drinking in rodents (36) through its anxiolytic and anti-stress properties, as CRF is the major regulator of the stress response (59). Dysregulation of the stress-related CRF circuitry by repeated alcohol use and induction of dependence leads to heightened anxiety and increased alcohol self-administration during withdrawal or thus negative reinforcement (60). Clinically, R121919 has shown efficacy in alleviating anxious and depressive symptoms in a human trial, but development was discontinued due to elevated liver enzyme activity in some participants (61). Development of safer CRHR1 antagonist alternatives, like Verucerfont, showed attenuated amygdala response to negative affective stimuli in patients with alcohol use disorder, but failed to reproduce the desired effect on alcohol craving (62). In the present study, R129191 treatment decentralized the CEA in the withdrawal network, resulting in reduced functional connectivity within the addiction network, with disinhibition of the frontal cortex. These findings align with the hypothesis that blocking CRHR1, primarily expressed in the cortical plate, normalizes the hyperactivity of the brain stress system originating from the CEA through GABAergic CRF neurons during withdrawal.

Naltrexone, an FDA-approved MOR antagonist for alcohol use disorder, reduces heavy drinking and craving (63) by decreasing dopaminergic activity and thus the positive reinforcement or rewarding effects of alcohol if drinking occurs (64, 65). In contrast to R121919, naltrexone reduced FOS expression and reactivity, increased brain-wide correlations, decreased modularity, and significantly enhanced the functional connectivity between prefrontal cortical and subcortical regions. Increased frontostriatal connectivity has also been observed in human fMRI studies, where it has been hypothesized to align with increased executive control over drinking and reduce wanting (66, 67). Debate exists in the literature regarding the necessity of abstinence before initiating naltrexone treatment (63, 68, 69) and reports of malaise, aversion, and naltrexone-induced side effects in patients, causing low patient compliance, but also correlating with treatment efficacy (70-73).Therefore, it is possible that the lack in normalization, but rather a further decrease in modularity following naltrexone treatment in dependent animals captured a dysphoric state, possibly due to precipitated alcohol withdrawal from MOR antagonism. Alcohol consumption affects the endogenous opioid system (74), which is inhibited by naltrexone. Indeed, we have unpublished data suggesting that while naltrexone indeed reduced drinking and hyperalgesia during withdrawal in alcohol-dependent rats, it also caused anhedonia. The observed hypomodular network may reflect this anhedonic state. Additionally, the upregulation of FOS expression in the lateral habenula (LH), which is rich in MOR, following naltrexone treatment in dependent mice, though not significant in our current sample, also pointed in this direction as the LH is considered the brain’s anti-reward center (75). Hence, the hypothesis of using modularity to screen for alcohol withdrawal and dependence may hold, considering that some treatments act through an anhedonic state.

A limitation of this study stems from the timing of the treatment in acute withdrawal, which could introduce confounding withdrawal precipitation effects. In the future, the treatments should also be tested later in abstinence. Moreover, a chronic treatment with naltrexone as advised for patients with alcohol use disorder should also be evaluated in addition to the current study of acute single injections.

Aside from the timeline, there are other noteworthy considerations and limitations. Firstly, the CIE model that is paired with two bottle choice (2BC) drinking is preferred for induction of dependence over the passive method of i.p. injections used here. A follow-up study comparing the effect on the functional connectivity of voluntary drinking, could explain why we do not see the strong anticorrelation with the amygdala as previously observed (1). Alternatively the difference can be related to the earlier withdrawal timepoint we are looking at (2). Additional and more complex behavioral measures like 2BC drinking would also improve our regression models. Secondly, while the study was well-powered and performed in both male and female subjects, the subdivision by sex (N=4M + 4F/group) lacks the power to represent the data here. Additionally, two female mice from the naltrexone-treated dependent group were lost, one during alcohol treatment and one during brain processing, compromising the correlation analysis in this subgroup. Preliminary indications suggest that the R121919 effect may be more pronounced in males than females (data not shown), aligning with observed sex differences in midbrain response to CRHR1 antagonists (76) and possibly contributing to the lack of CRHR1 antagonist efficacy in clinical trials which were performed in females (62). This underscores the importance of including male and female subjects in research studies and the need for follow-up studies to increase the power to detect sex differences. Even with FDA approval, the usefulness of naltrexone in women is understudied (77). Lastly, a more general note pertains to relying on FOS expression as a marker of neuronal reactivity, recognizing that it may not capture all relevant aspects of neuronal activity. c-Fos was selected over other immediate early genes because its low baseline expression levels, optimal for detecting increases in neuronal activity, past success with the iDISCO+ technique (14, 17), and our prior data using regular slide FOS immunohistochemistry. Future investigations will explore additional immediate early genes and co-label neuronal markers to comprehensively study cell-type-specific responses.

Despite these limitations, future plans, and challenges, the current study revealed significant changes in the withdrawal and control networks following treatment with R129191 and naltrexone. R121919, acting on the CRHR1 receptors, partially normalized modularity by activating the prefrontal cortex and reducing the influence of the extended amygdala on the network, effectively blocking negative reinforcement through the stress system. Conversely, naltrexone, acting on the MORs, inhibited most basal forebrain, amygdala, and hypothalamic regions, increasing functional connectivity between the prefrontal cortex and subcortical areas to block positive reinforcement through the endogenous opioid system. The functional connectomic approach can thus provide crucial insights into and help understand treatments for alcohol use disorder, which is essential for advancing the development of novel therapeutic interventions.

## Acknowledgments

Support from the Preclinical Research Addiction Consortium is acknowledged. This work was supported by funding from NIAAA P60AA006420 to OG and an Innovation Award from the International Rett Syndrome Foundation to LC. Light-sheet microscopy was performed at the California Institute of Technology (Caltech), Biological Imaging Facility with the support of the Caltech Beckman Institute and the Arnold and Mabel Beckman Foundation. Publication fees were contributed by the UCSD library.

## Notes

### Competing Interest Statement

The authors have declared no competing interest.

### Summary of Updates

Correct the heatmap of figure 4A in which the rows where not correctly ordered.

