## Extended data for "Antagonists of the stress and opioid systems restore the functional connectivity of the prefrontal cortex during alcohol withdrawal through divergent mechanisms"

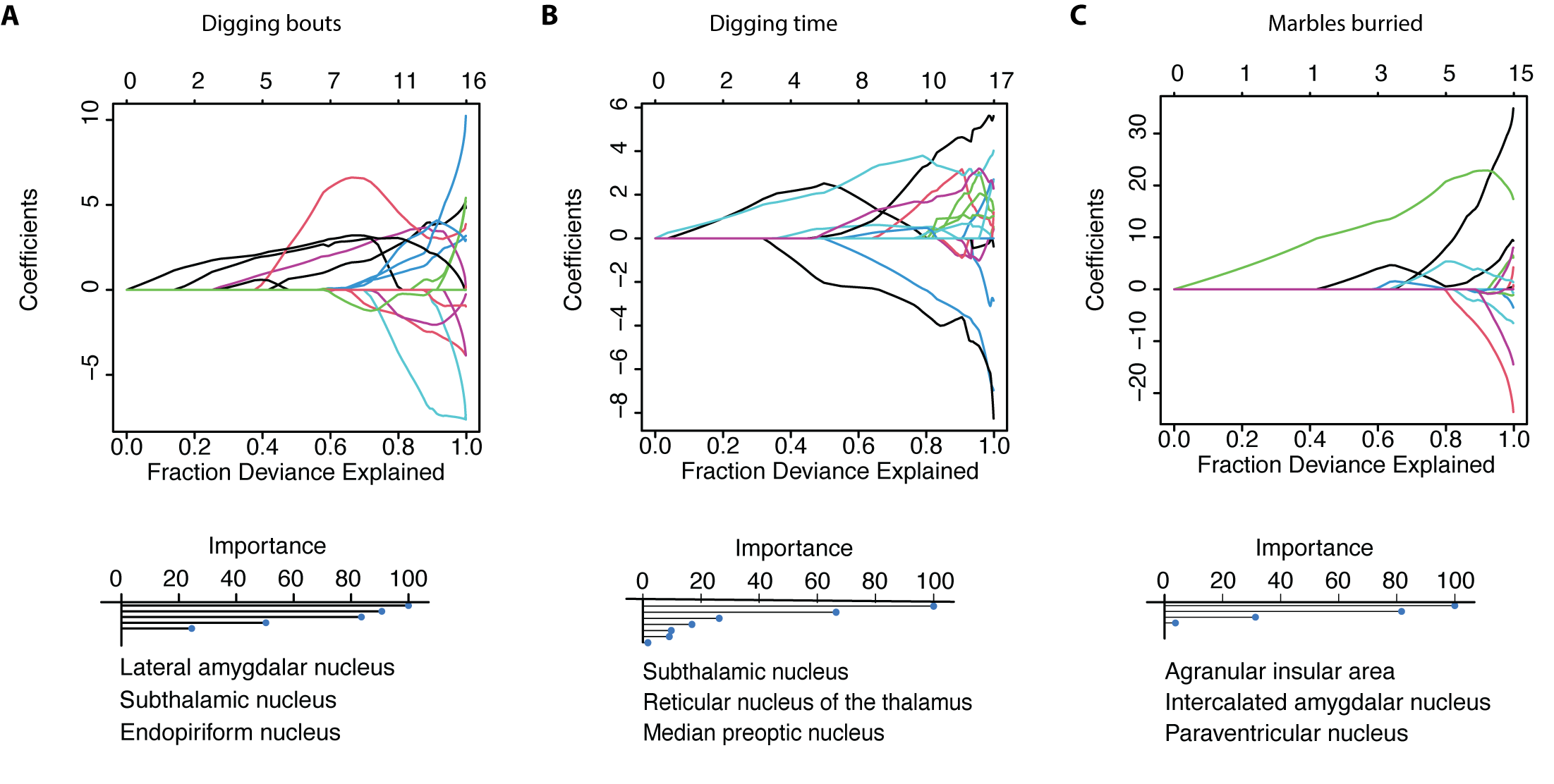

Figure S1: *LASSO regression model fitting of all saline treated animals (dependent and non-dependent), showing the number of variables (top x-axis), their coefficients (y-axis) in function of the fraction of deviance explained (bottom x-axis), and the most important identified variables or brain regions of the model for bouts of digging (A), length of digging (B), and number of marbles buried (C). To avoid the introduction of any variation that was not captured in the behavior as noise, only the saline treated control and dependent animals were included in the analysis.*

The lateral amygdalar nucleus (LA), subthalamic nucleus (STN), and endopiriform nucleus (EP) were the top regions for bouts of digging (Fig. S1A). The STN was also a top regions for length of digging together with the reticular nucleus of the thalamus (RT), and median preoptic nucleus (MPN) (Fig. S1B). For the number of marbles buried, which was not significantly different between dependent animals in withdrawal and their controls, the agranular insular area (AI), intercalated amygdalar nucleus (IA) and paraventricular nucleus of the thalamus (PVT) were the top regions identified (Fig. S1C).

Several of the identified main contributing regions to the LASSO model had previously been identified to have important roles in alcohol dependency. The IA, LA and EP, were found previously to cluster together in the withdrawal network (1). The STN that is a common contributor to both the digging time and bouts models has previously been shown to have upregulated FOS activity during withdrawal in rodents (2) and to have a different connectivity in patient with alcohol use disorder compared to healthy controls (2). The marble burying model, which did not show a significant difference between dependent and control animals when counting the amount of marbles, also identified regions important for alcohol withdrawal, the AI, and PVT. The AI was found to increase activity during withdrawal from chronic alcohol exposure in rats (3) and its reduced functional connectivity has been associated with reduced alcohol drinking in patients with alcohol use disorder (4). The PVT is a key anxiety regulator that integrates cortical and hypothalamic signals to modulate behavior (5), lesions have been shown to prevent alcohol relapse (6)

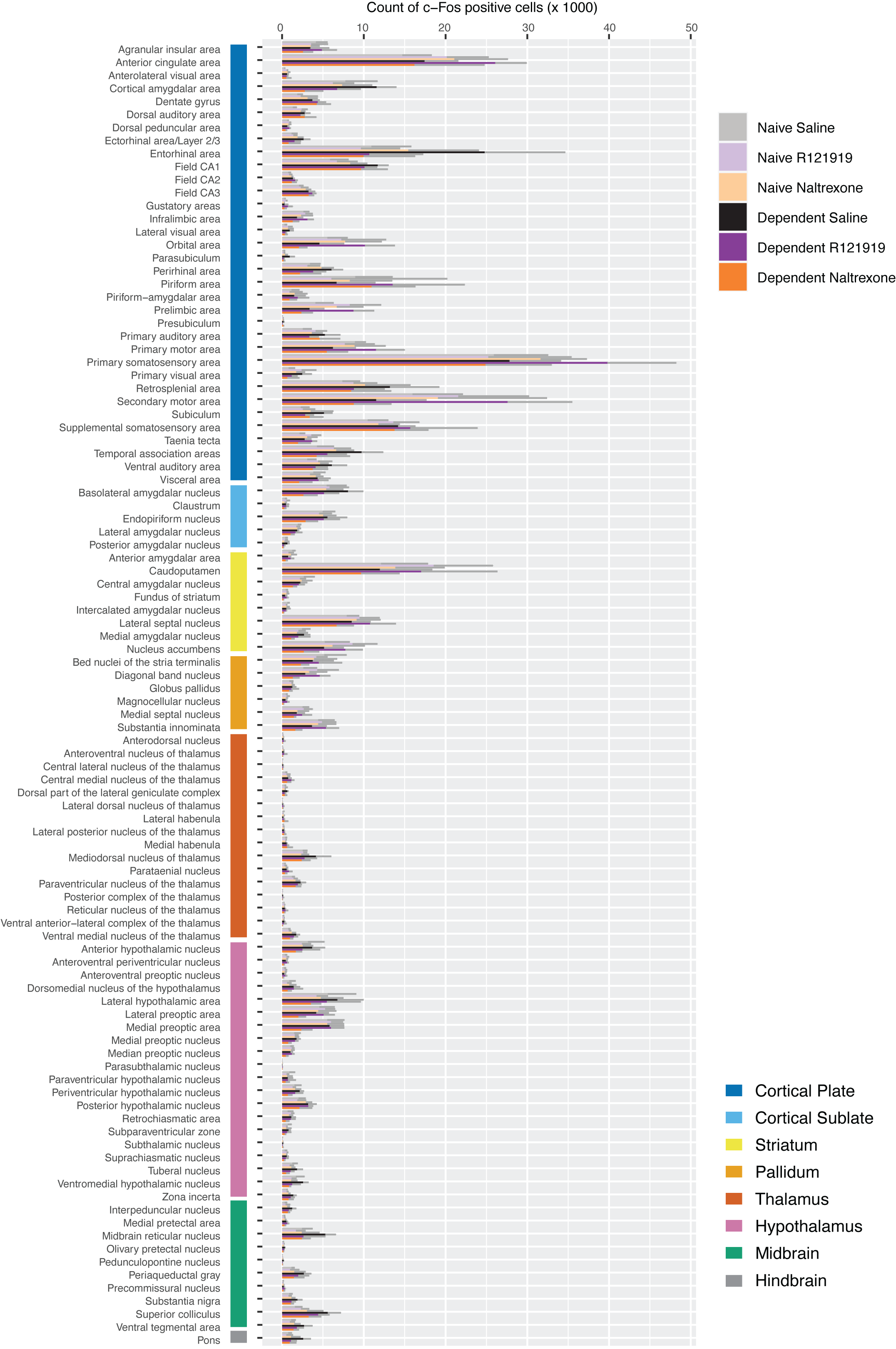

*Figure S2. Supplement for Figure 2, raw Fos counts per region*

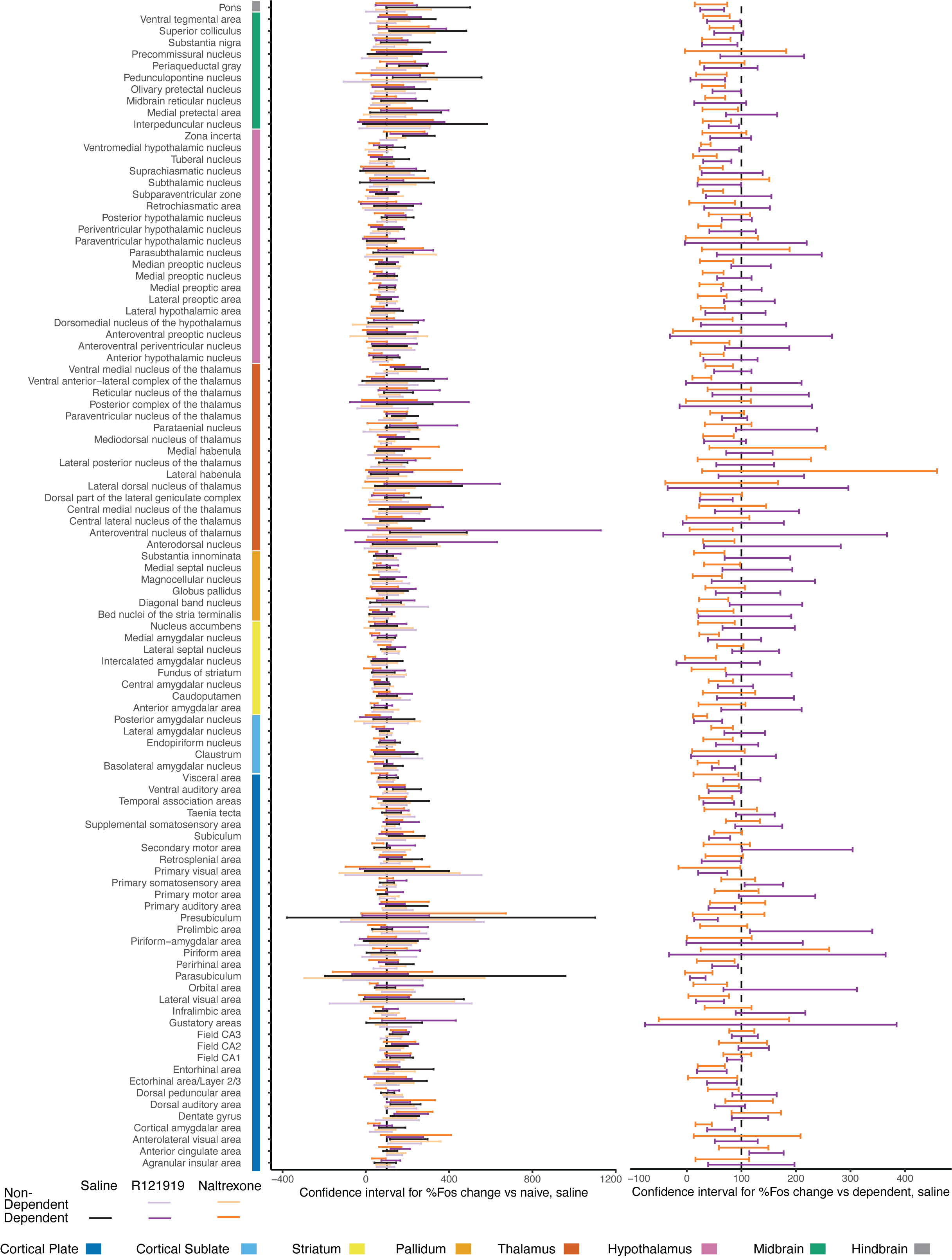

Figure S3. Supplement for Fig. 2, **all 95%-confidence intervals.**

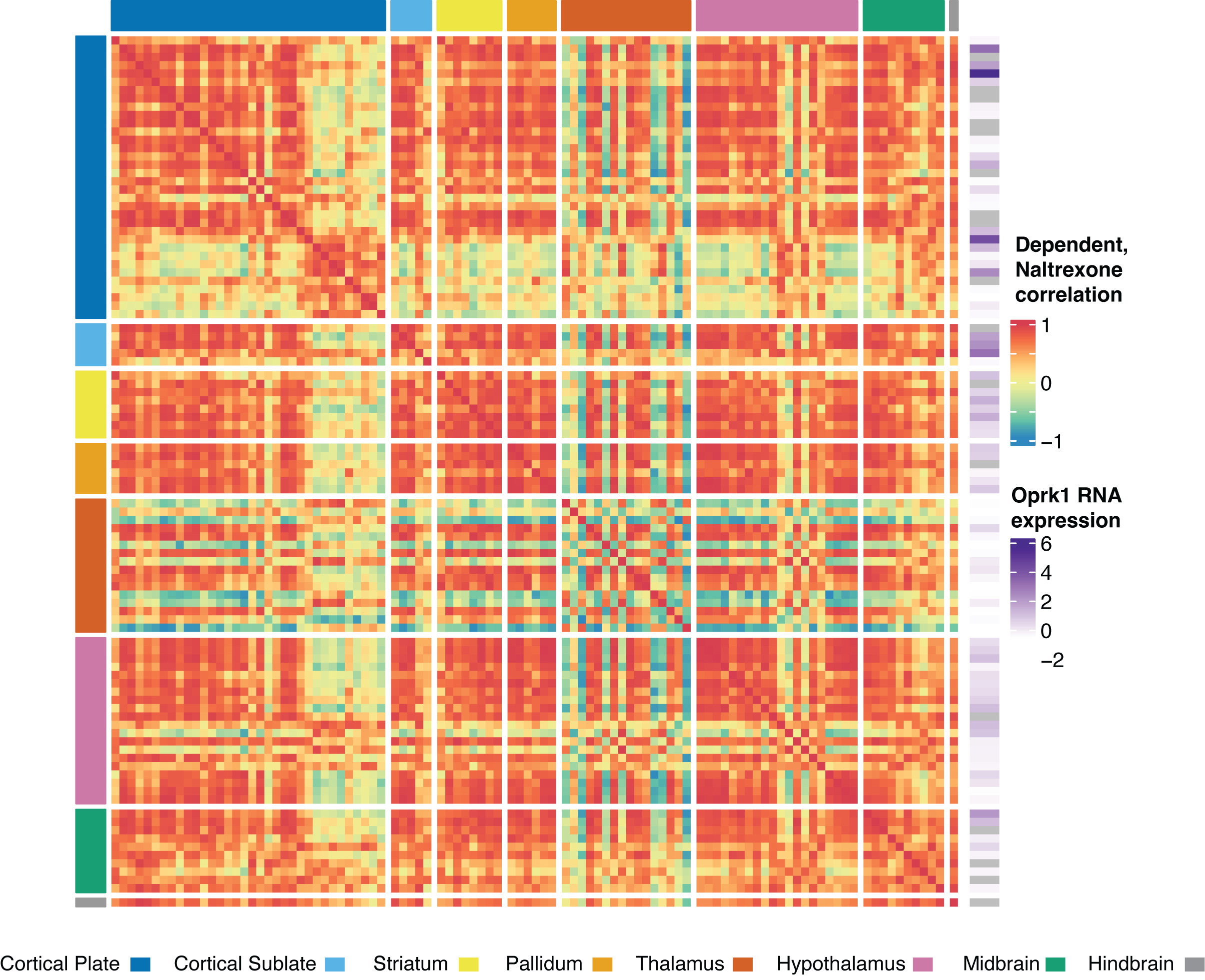

*Figure S4. Supplement for Figure 4,* ***Oprk1 mRNA expression (purple) combined with whole-brain functional connectomics for alcohol dependent Naltrexone treated animals.***

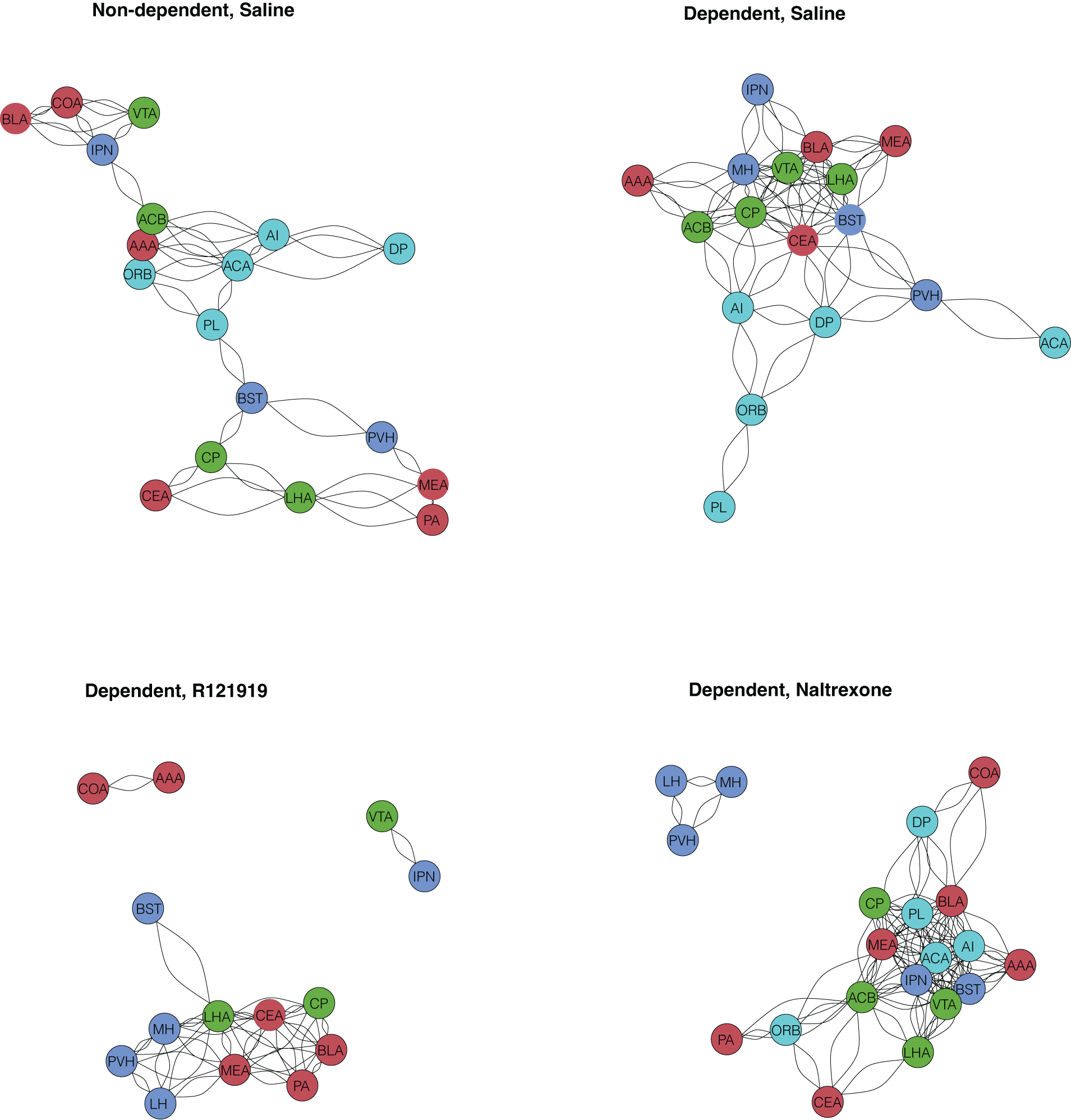

*Figure S5. Supplemental data for Figure 5,* ***The minimal addiction networks represented following Force Field simulation.***

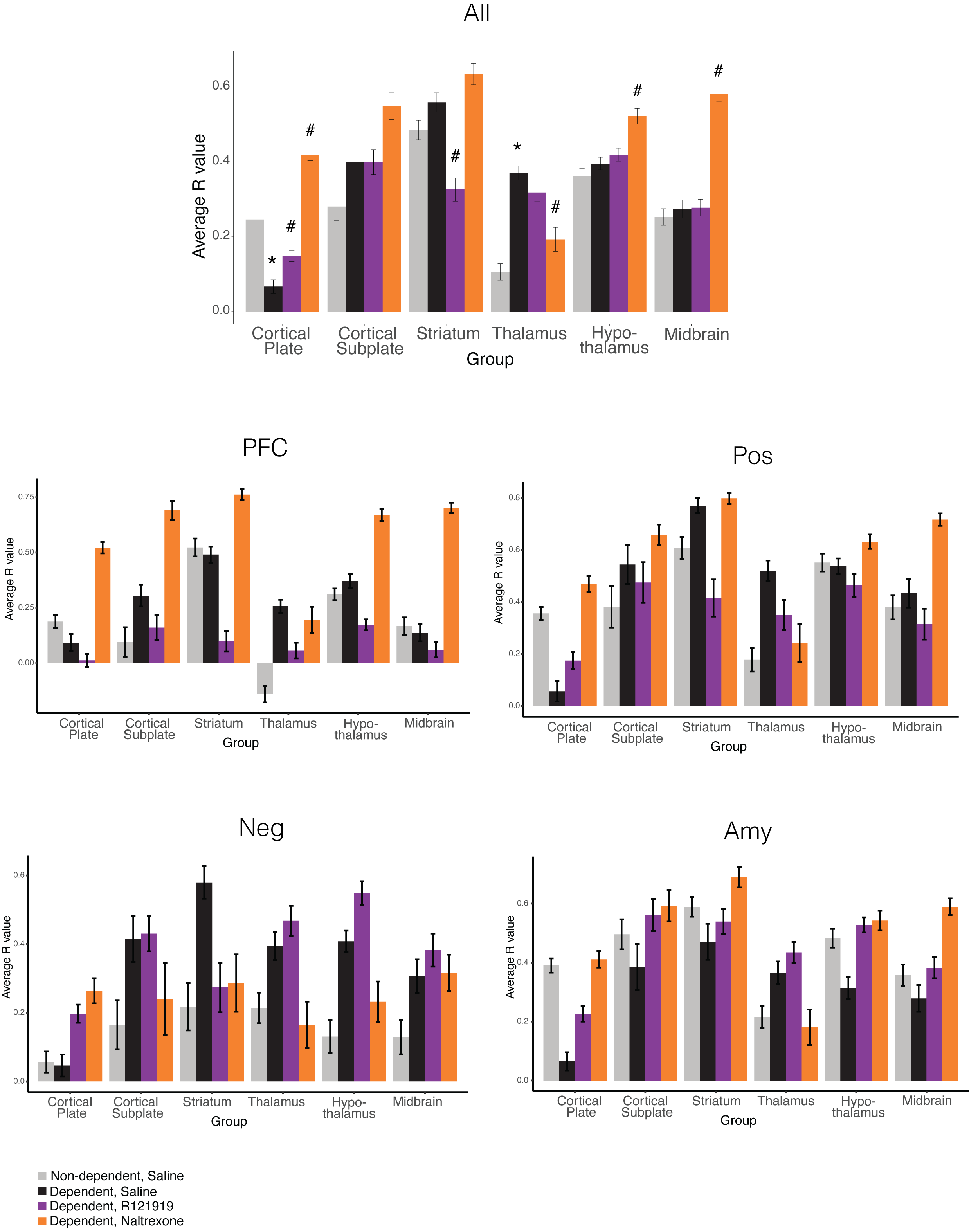

*Figure S6. Supplement to Figure 5,* ***Average correlations of the addiction network with the rest of the brain*** *per anatomical group for the different treatment groups, for the whole addiction sub-network (*vs non-dependent saline, ^#^ vs dependent saline) or split for regions in the frontal cortex (PFC), associated with positive (Pos) and negative (Neg) emotional state, and the amygdala (Amy).*

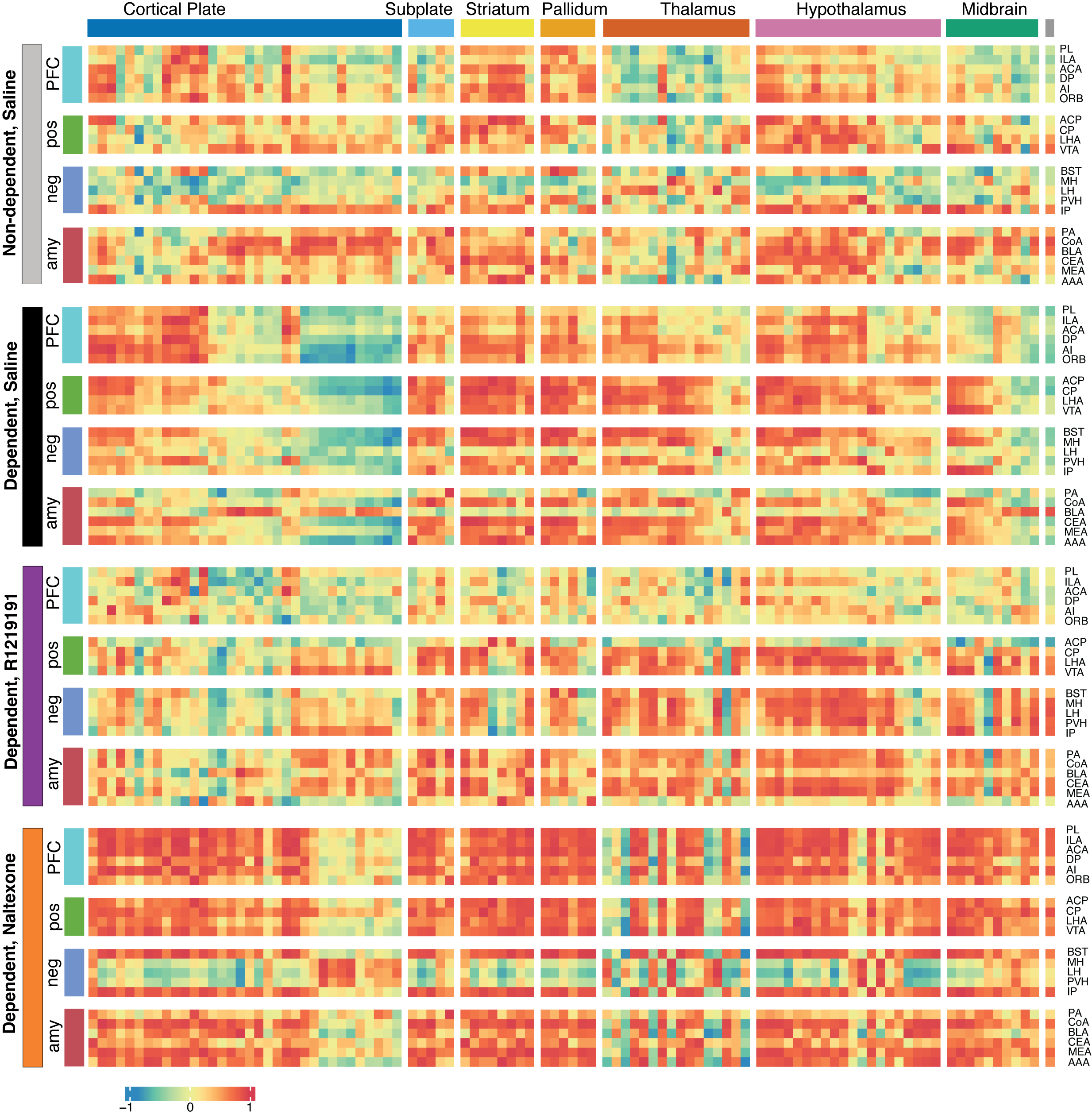

*Figure S7. Supplement to Figure 5,* ***Correlation of the regions of the addiction network with the rest of the brain for the different treatment groups.*** *Heatmap of correlations of all brain regions with the addiction network, consisting of regions in the frontal cortex (turquoise), associated with positive (green) and negative (blue) emotional state, and the amygdala (red). Brain regions are organized according to the hierarchical clustering of the saline-treated dependent animals per anatomical group (as in Fig. 4 and Table S2).*

Table S1: Alphabetical organized list of brain regions with abbreviations

| region | abbreviation | group |
| --- | --- | --- |
| Agranular insular area | AI | Cortical Plate |
| Anterior amygdalar area | AAA | Striatum |
| Anterior cingulate area | ACA | Cortical Plate |
| Anterior hypothalamic nucleus | AHN | Hypothalamus |
| Anterodorsal nucleus | AD | Thalamus |
| Anterolateral visual area | VISal | Cortical Plate |
| Anteroventral nucleus of thalamus | AV | Thalamus |
| Anteroventral periventricular nucleus | AVPV | Hypothalamus |
| Anteroventral preoptic nucleus | AVP | Hypothalamus |
| Basolateral amygdalar nucleus | BLA | Cortical Subplate |
| Bed nuclei of the stria terminalis | BST | Pallidum |
| Caudoputamen | CP | Striatum |
| Central amygdalar nucleus | CEA | Striatum |
| Central lateral nucleus of the thalamus | CL | Thalamus |
| Central medial nucleus of the thalamus | CM | Thalamus |
| Claustrum | CLA | Cortical Subplate |
| Cortical amygdalar area | COA | Cortical Plate |
| Dentate gyrus | DG | Cortical Plate |
| Diagonal band nucleus | NDB | Pallidum |
| Dorsal auditory area | AUDd | Cortical Plate |
| Dorsal part of the lateral geniculate complex | LGd | Thalamus |
| Dorsal peduncular area | DP | Cortical Plate |
| Dorsomedial nucleus of the hypothalamus | DMH | Hypothalamus |
| Ectorhinal area/Layer 2/3 | ECT | Cortical Plate |
| Endopiriform nucleus | EP | Cortical Subplate |
| Entorhinal area | ENT | Cortical Plate |
| Field CA1 | CA1 | Cortical Plate |
| Field CA2 | CA2 | Cortical Plate |
| Field CA3 | CA3 | Cortical Plate |
| Fundus of striatum | FS | Striatum |
| Globus pallidus | PALd | Pallidum |
| Gustatory areas | GU | Cortical Plate |
| Infralimbic area | ILA | Cortical Plate |
| Intercalated amygdalar nucleus | IA | Striatum |
| Interpeduncular nucleus | IPN | Midbrain |
| Lateral amygdalar nucleus | LA | Cortical Subplate |
| Lateral dorsal nucleus of thalamus | LD | Thalamus |
| Lateral habenula | LH | Thalamus |
| Lateral hypothalamic area | LHA | Hypothalamus |
| Lateral posterior nucleus of the thalamus | LP | Thalamus |
| Lateral preoptic area | LPO | Hypothalamus |
| Lateral septal nucleus | LSX | Striatum |
| Lateral visual area | VISl | Cortical Plate |
| Magnocellular nucleus | MA | Pallidum |
| Medial amygdalar nucleus | MEA | Striatum |
| Medial habenula | MH | Thalamus |
| Medial preoptic area | MPO | Hypothalamus |
| Medial preoptic nucleus | MPN | Hypothalamus |
| Medial pretectal area | MPT | Midbrain |
| Medial septal nucleus | MS | Pallidum |
| Median preoptic nucleus | MEPO | Hypothalamus |
| Mediodorsal nucleus of thalamus | MD | Thalamus |
| Midbrain reticular nucleus | MRN | Midbrain |
| Nucleus accumbens | ACB | Striatum |
| Olivary pretectal nucleus | OP | Midbrain |
| Orbital area | ORB | Cortical Plate |
| Parasubiculum | PAR | Cortical Plate |
| Parasubthalamic nucleus | PSTN | Hypothalamus |
| Parataenial nucleus | PT | Thalamus |
| Paraventricular hypothalamic nucleus | PVH | Hypothalamus |
| Paraventricular nucleus of the thalamus | PVT | Thalamus |
| Pedunculopontine nucleus | PPN | Midbrain |
| Periaqueductal gray | PAG | Midbrain |
| Perirhinal area | PERI | Cortical Plate |
| Periventricular hypothalamic nucleus | PV | Hypothalamus |
| Piriform area | PIR | Cortical Plate |
| Piriform-amygdalar area | PAA | Cortical Plate |
| Pons | P | Hindbrain |
| Posterior amygdalar nucleus | PA | Cortical Subplate |
| Posterior complex of the thalamus | PO | Thalamus |
| Posterior hypothalamic nucleus | PH | Hypothalamus |
| Precommissural nucleus | PRC | Midbrain |
| Prelimbic area | PL | Cortical Plate |
| Presubiculum | PRE | Cortical Plate |
| Primary auditory area | AUDp | Cortical Plate |
| Primary motor area | MOp | Cortical Plate |
| Primary somatosensory area | SSp | Cortical Plate |
| Primary visual area | VISp | Cortical Plate |
| Reticular nucleus of the thalamus | RT | Thalamus |
| Retrochiasmatic area | RCH | Hypothalamus |
| Retrosplenial area | RSP | Cortical Plate |
| Secondary motor area | MOs | Cortical Plate |
| Subiculum | SUB | Cortical Plate |
| Subparaventricular zone | SBPV | Hypothalamus |
| Substantia innominata | SI | Pallidum |
| Substantia nigra | SN | Midbrain |
| Subthalamic nucleus | STN | Hypothalamus |
| Superior colliculus | SC | Midbrain |
| Supplemental somatosensory area | SSs | Cortical Plate |
| Suprachiasmatic nucleus | SCH | Hypothalamus |
| Taenia tecta | TT | Cortical Plate |
| Temporal association areas | TEa | Cortical Plate |
| Tuberal nucleus | TU | Hypothalamus |
| Ventral anterior-lateral complex of the thalamus | VAL | Thalamus |
| Ventral auditory area | AUDv | Cortical Plate |
| Ventral medial nucleus of the thalamus | VM | Thalamus |
| Ventral tegmental area | VTA | Midbrain |
| Ventromedial hypothalamic nucleus | VMH | Hypothalamus |
| Visceral area | VISC | Cortical Plate |
| Zona incerta | ZI | Hypothalamus |

Table S2: order of the regions following hierarchical clustering and separation into modules

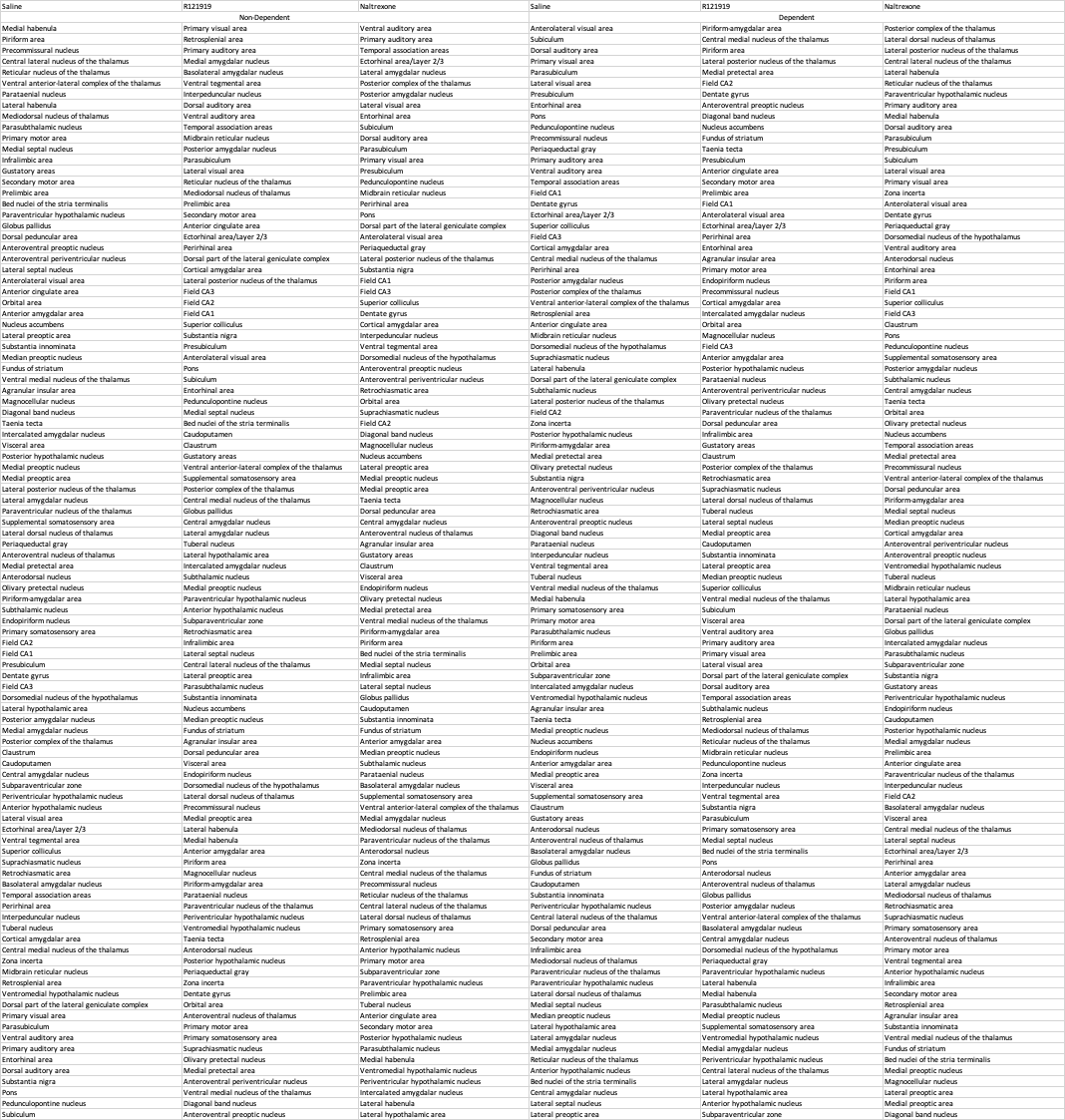

Table S3: Organization of the brain regions per anatomical group and clustered according to saline treated dependent animals

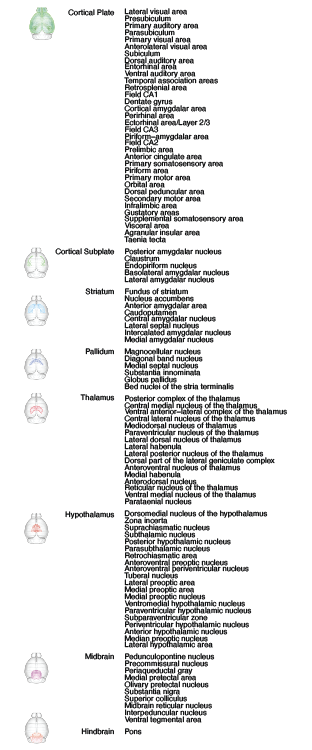

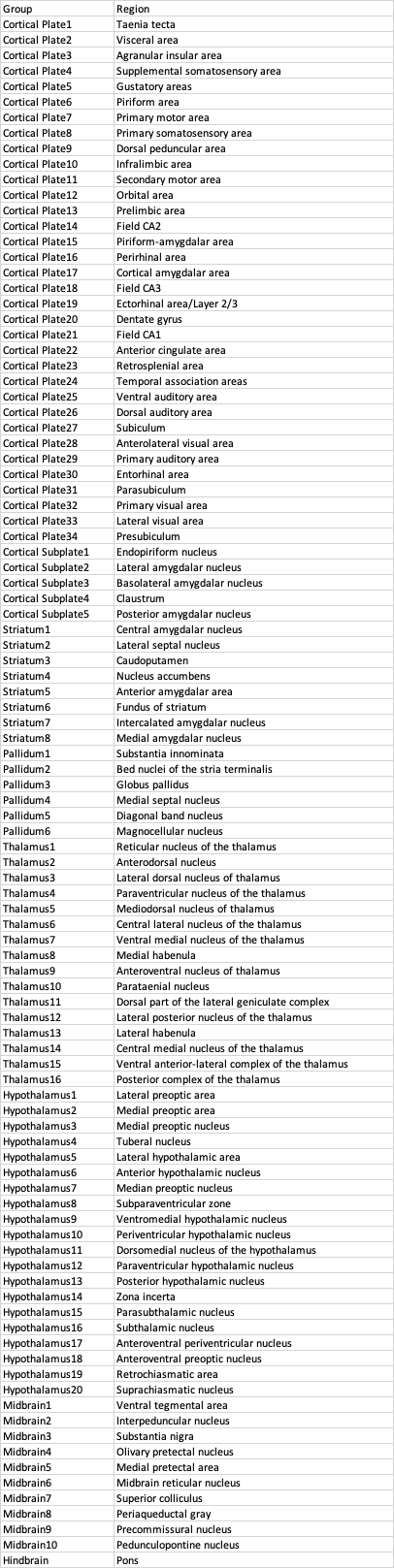
